# Elucidating Direct Kinase Targets of Compound Danshen Dropping Pills Employing Archived Data and Prediction Models

**DOI:** 10.1101/2020.03.13.990291

**Authors:** Tongxing Wang, Lu Liang, Chunlai Zhao, Jia Sun, Hairong Wang, Wenjia Wang, Jianping Lin, Yunhui Hu

**Affiliations:** GeneNet Pharmaceuticals Co. Ltd., No.1, Tingjiang West Road, Beichen District, Tianjin 300410, China; College of Pharmacy, Nankai University, 38 Tongyan Road, Haihe Education Park, Jinnan District, Tianjin 300353, China

**Keywords:** Compound Danshen Dropping Pill, Traditional Chinese Medicine, Kinase, Direct target, Allosteric effect, Mechanism of action

## Abstract

The research on the direct target of traditional Chinese medicine (TCM) is the key to study the mechanism and material basis of TCM, but there is still no effective technical methods at present. For Compound Danshen dropping pills (CDDP), there is no report about its direct targets. In this study, the direct targets of CDDP were studied for the first time, especially focusing on the protein kinase family, which plays causal roles in a variety of human disease. Firstly, the literature database of CDDP was constructed by literature retrieval, and the important components contained in CDDP were extracted. Secondly, the potential direct targets of important components was obtained through querying public database and predicted by Multi-voting SEA algorithm. Then, the KinomeX system was used to predict and to filter the potential kinase targets of CDDP. Finally, the experimental verification was carried out. In total, 30 active kinase targets was obtained at 25 μg/ml concentration of CDDP, and 9 dose-dependent targets were obtained at 250 μg/ml concentration of CDDP. This is an efficient and accurate strategy by integrating the targets recorded in several public databases and the targets calculated by two in silico modelling approaches predict potential direct targets of TCM, which can lay an important foundation for the study of the mechanism and material basis of them, promoting the modernization of TCM.

## Introduction

Traditional Chinese medicine prescriptions are the characteristics of TCM. They have been practiced for thousands of years and have been proved to be effective in modern clinical practice. These prescriptions embodies the dialectical thought of TCM and the medication holistic view. However, there is still a lack of effective approaches to systematically study its mechanism and material basis. In recent years, the reductionist research model has accumulated a lot of data, and also provided illuminating research results, such as the discovery of artemisinin [1]. However, considering the complexity of TCM, it is still difficult to completely separate and identify the effective components through the existed technology and analytical methods, let alone to make clear all the effective components in the TCM. Therefore, the research of reductionism is not capable of answering the essential question of the overall efficacy of TCM. It may lead to deviate from the system theory of TCM, so it needs to be combined with the system theory. In recent years, a variety of “omics” techniques based on system theory have been widely used in the research of TCM [2–5], which can be used to better understand the pharmacological characteristics of TCM [6–9], but still cannot fully reveal the nature of it. Comprehensively understanding the mechanism of synergism among the effective components, drug targets and metabolic pathways is still highly demanded. One key to solve the problem is to carry out the research on the direct target of TCM. It can not only clarify the pharmacological mechanism of TCM from the origin and scientifically interpret its traditional efficacy, but also unveil new disease mechanism [10] and provide reasonable estimation of TCM repositioning. However, many obstacles cannot be ignored in the study of the direct target of TCM, such as the complex ingredients in it, the complex process of metabolism in vivo, the complicated way playing a role in vivo, and so on. At present, technical methods to screen and determine the direct targets for TCM efficiently and accurately are still poorly developed, which hinders elucidating the mechanisms of TCM essentially.

Compound Danshen dropping pills (CDDP) consist of *Radix Salviae* (Danshen), *Panax Notoginseng (Burk.) F. H. Chen Ex C. Chow* (Sanqi), *Borneolum Syntheticum* (Bingpian). It is widely used in the prevention, treatment and emergency treatment of coronary artery disease (CAD) and angina pectoris. Although many research articles about CDDP have been published already, the research on its mechanism of action is still not in-depth [11–14]. Most studies focused on the genes or proteins regulated by CDDP treatment, most of which can be referred as indirect targets, but there is no report on the direct targets of CDDP.

At present, kinases belong to an important class of drug targets. Among them, protein kinases family is the largest group of kinases, which act on specific proteins and change their activities. These kinases play a wide range of roles in cell signaling and complex life activities, and their dysfunction plays an important causal role in many human diseases, including cancer, inflammatory diseases, central nervous system diseases, cardiovascular diseases and so on [15].

In view of the importance of kinases, we proposes a new systematic approach to explore direct kinase targets of TCM efficiently and reliably, CDDP was taken as the research subject. This strategy is based on the known activity data recorded in free databases and predicted data calculated by computational models, which is independent of any specific disease model. Firstly, the literature database of CDDP was constructed by literature retrieval, and the important components contained in CDDP were extracted. Secondly, the potential targets of important components was obtained through public database querying and Multi-voting SEA algorithm predicting. Then, the KinomeX system was used to filter the potential kinase targets of CDDP. Finally, 30 active targets were obtained (Figure 1). The activity of MET, PIM1 and SYK had obvious dose-dependent effect, and that of CAMK2G, CSF1R, FYN and RET increased at high concentration, which are worthy of further study.

**Figure 1.**
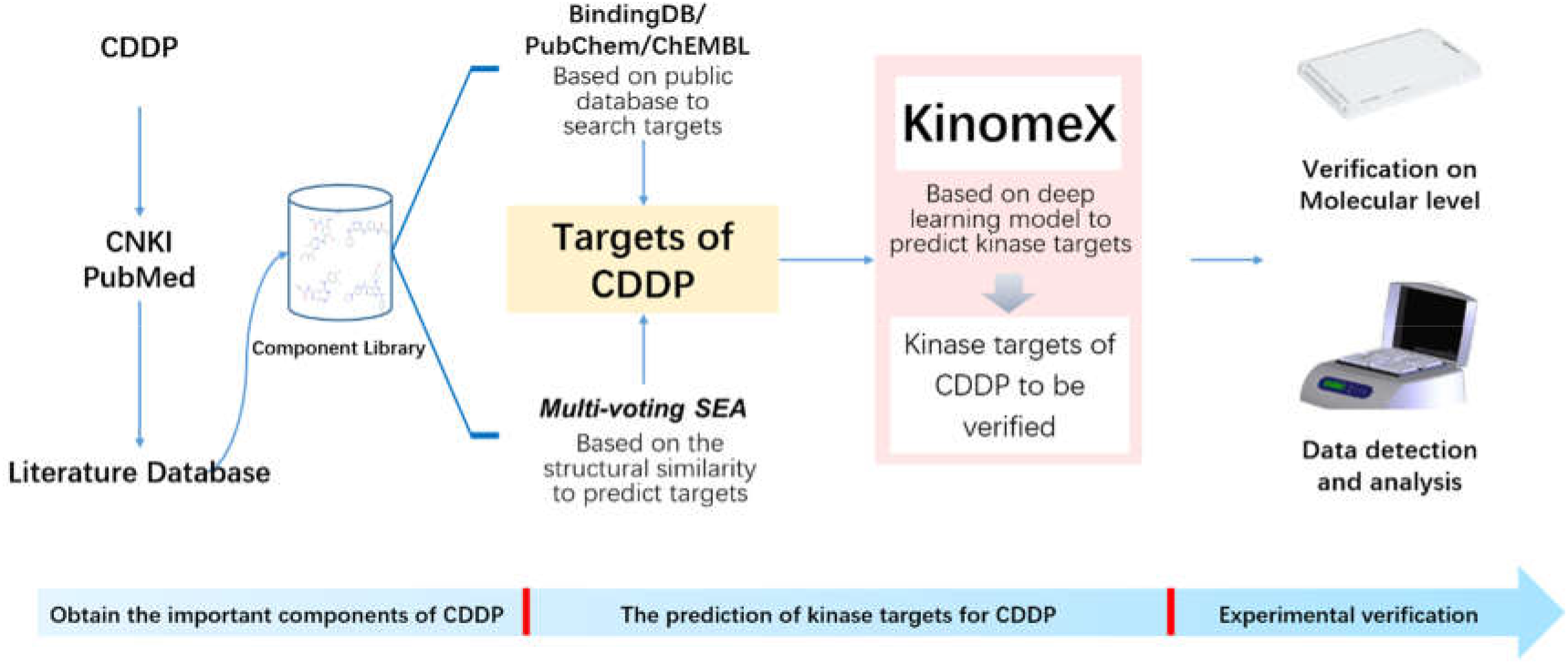
The flowchart of the study based on known activity data and prediction model to obtain the direct kinase targets of CDDP

## Methods

### 1 Construction of important component set for CDDP

In order to review the literatures related to CDDP as comprehensively as possible, we firstly customized the relevant search terms of it. The Chinese literature retrieval word was “Danshen Dropping Pills”, and the English literature retrieval words were “Compound Danshen dripping pills”, “Fufang Danshen Diwan”, “T89”, “Dantonic” and “Cardiotonic Pills”. Secondly, we used “Danshen Dropping Pills” as the keyword to obtain the Chinese-language literatures through CNKI (https://www.cnki.net/). Similarly, through PubMed (https://pubmed.ncbi.nlm.nih.gov/), “Compound Danshen drilling pills”, “Fufang Danshen Diwan”, “T89”, “dantonic” and “Cardiotonic Pills” were used to get the English-language literatures (time to December 15, 2019). Finally, the literature database of CDDP was constructed by combining the above literatures from two sources.

The components contained in CDDP were extracted manually by reading the literature, and then the information of them was standardized through PubChem database (https://pubchem.ncbi.nlm.nih.gov/) [16]. It is generally believed that the ingredients entering the blood, main metabolites, bioequivalence components compared to the prescription, and active components of CDDP reported in literatures are the important components in CDDP. Additionally, in order to avoid missing critical components included in CDDP, we selected the most extensively studied component in the three single herbs but still unconfirmed in the whole prescription, through retrieving TCM related databases, such as TcmSP [17]、TCMID [18]、TCM-ID [19]、ETCM [20]、YaTCM [21].

### 2 Prediction of kinases targets of CDDP

The known activity data of 40 important components in CDDP were obtained from three authoritative public databases, namely, ChEMBL [22], PubChem [16], BingdingDB [23]. The kinase targets with definite activity information were standardized and screened.

Avoiding missing some important potential kinase targets, Multi-voting SEA algorithm [24] was utilized to predict potential direct targets of important components. In this algorithm, four scoring models, namely Topological SEA, Morgan SEA, MACCS SEA Atom Pair SEA, were integrated to calculate potential targets of components. Finally, all potential targets of each component were normalized and the kinase targets were selected.

The KinomeX system (https://kinome.dddc.ac.cn/en/) [25] developed by Shanghai Pharmaceutical Institute was used to predict the potential kinase targets of 40 important components of CDDP. It enables users to predict its potential kinase targets based on the structure of a given molecule. The prediction results were used to screen the targets obtained from public databases and Multi-voting SEA algorithm mentioned above. The kinase targets screened by results from KinomeX were subsequently taken as the kinase target set of CDDP with high reliability to conduct following experimental verification.

### 3 Experimental verification for direct targets of CDDP

Full KP panel [km ATP], a kinaseprofiler, was developed by Eurofins company. In this study, we used this panel to carry out experimental verification for direct kinase targets of CDDP. Firstly, the filter binding radioactive kinase activity assays were performed by using 25 μg/ml of CDDP. The kinase activity inhibition rate of the sample were expressed as the percentage of the result of sample compared to the blank group. The kinase activity of the blank was considered to be 100%. Generally speaking, if the residual enzyme activity is less than 30%, it is considered to be strongly inhibited. And if the residual enzyme activity is between 30% and 70%, it is considered as moderate inhibition. Considering the weak interaction superposition characteristic and synergistic effect of TCM ingredients [26, 27], the threshold value in this study was set to 80%.

In order to get the dose-dependent kinase targets, the kinase targets with activity value less than 70 were retested at 250 μg/ml of CDDP. The targets with obvious dose dependence characteristic were selected to get the IC50 curve through multi-concentration test.

## Results

### 1 Important components in CDDP

3719 Chinese-language literatures and 59 English-language literatures were obtained through retrieving the customized terms of the CDDP (time to December 15, 2019). Through literature reading manually, the components information of CDDP was extracted. According to the screening criteria of important ingredients, a total of 39 ingredients were collected. In addition, quercetin, a potential important component of the whole prescription was also included for subsequent analysis, which was reported a lot in single herbs but has not been confirmed in the whole prescription. All 40 important components of CDDP are shown in Table 1.

**Table 1.**
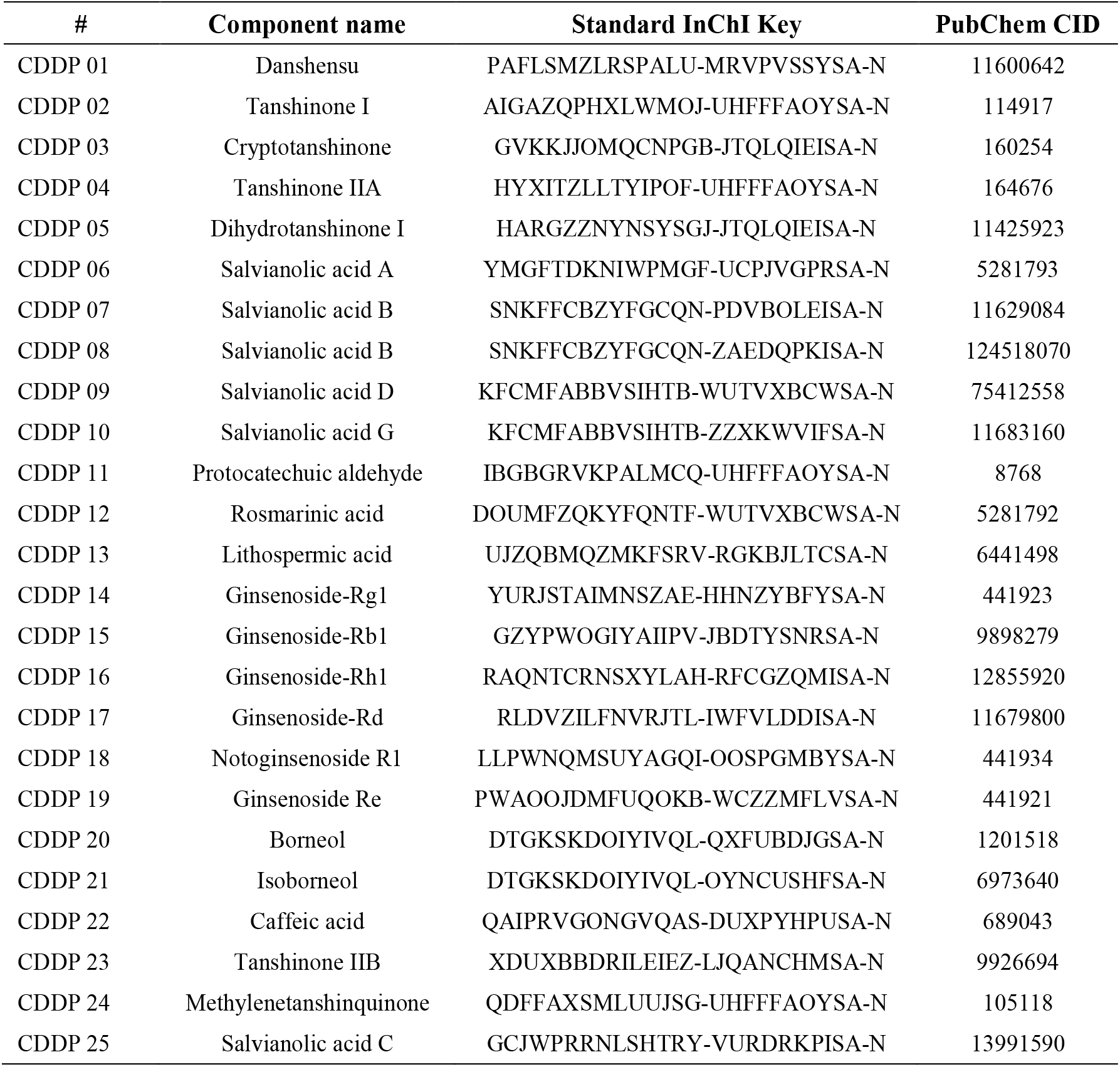
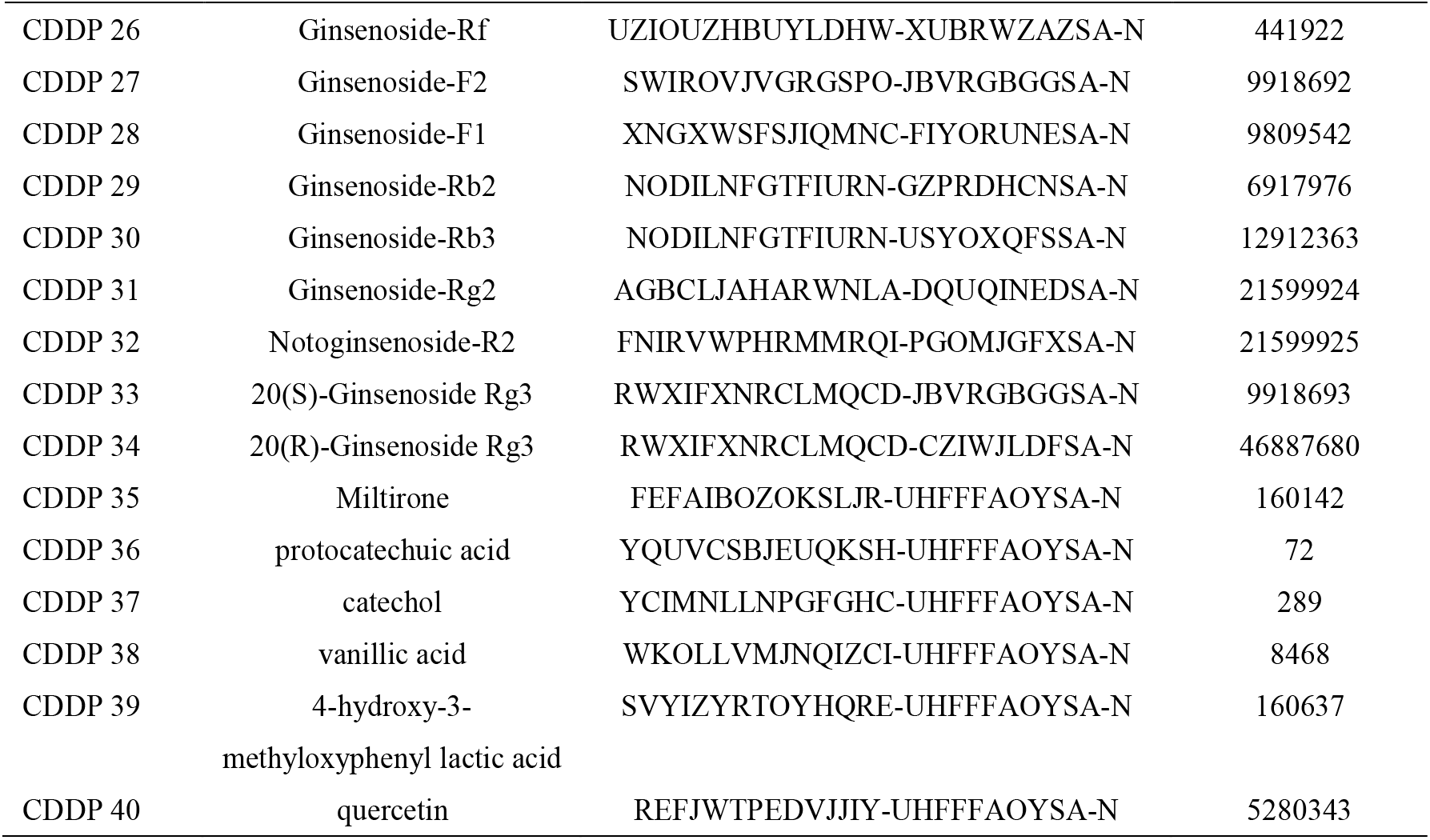
40 important components of CDDP

### 2 The potential kinase targets of CDDP

Based on the hypothesis that the potential direct targets of important components in the whole prescription are more likely to be direct targets of the whole prescription we carried out in the following analysis. Therefore, the potential targets of components were used to speculate the potential direct targets of the whole prescription. Three authoritative public databases and Multi-voting SEA algorithm were utilized to obtain direct targets information of important components. Moreover, the KinomeX system was used to predict the kinase targets, and finally the whole set of kinase targets of CDDP was obtained. Through querying the three public databases, 262 known targets including 55 kinase targets (Figure. 2, Table S1) were obtained, and 377 targets including 121 kinase targets (Figure. 2, Table S2) were predicted based on the Multi-voting SEA algorithm. By integrating the above two parts of targets, a total of 479 potential direct targets were obtained, including 148 kinase targets (Table 2, Table S3).

**Table 2.** Statistics of the number of potential direct targets for 40 important components of CDDP

| Target sources | Number of targets | Number of kinase targets |
| --- | --- | --- |
| Recorded targets | 262 | 55 |
| Multi-Voting SEA predicted targets | 377 | 121 |
| Common targets | 160 | 28 |
| Total targets | 479 | 148 |

**Figure 2.**
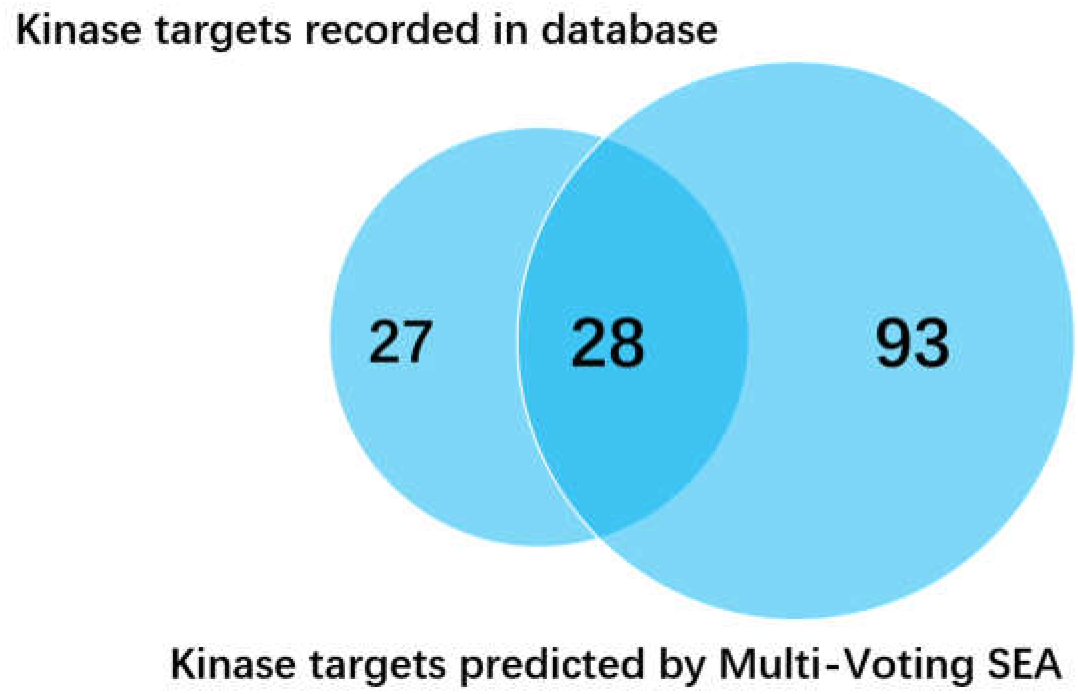
The number of potential direct kinase targets of CDDP

KinomeX, established by the research team of Shanghai Institute of pharmaceutical research, Chinese Academy of Sciences in 2019, is a prediction and analysis platform of single compound regulated kinase spectrum based on the existing big data of kinase activity and the deep neural network algorithm. Based on this method, the average ROC of this method is as high as 0.75, and the accuracy is significantly higher than other prediction methods [28–34]. Therefore, we used the KinomeX to predict the potential protein kinase targets of 40 important components in CDDP, and obtained 288 kinase targets (Table S4). And then we took this result to filter the 148 targets from above, and obtained the potential direct target set of CDDP for further verification. 37 kinase targets were selected from targets recorded in the databases (Table S5), and 92 kinase targets were selected from Multi-voting SEA predicted results (Table S6). These two parts shared 20 kinase targets (Figure 3), while 109 kinase targets deserve to be verified in total (Table 3, Table S7).

**Table 3.** Statistics of the number of potential direct kinase targets of CDDP to be verified

| Target sources | Number of targets | Number of kinase targets | Targets screened by KinomeX |
| --- | --- | --- | --- |
| Recorded targets | 262 | 55 | 37 |
| Multi-Voting SEA predicted targets | 377 | 121 | 92 |
| Common targets | 160 | 28 | 20 |
| Total targets | 479 | 148 | 109 |

**Figure 3.**
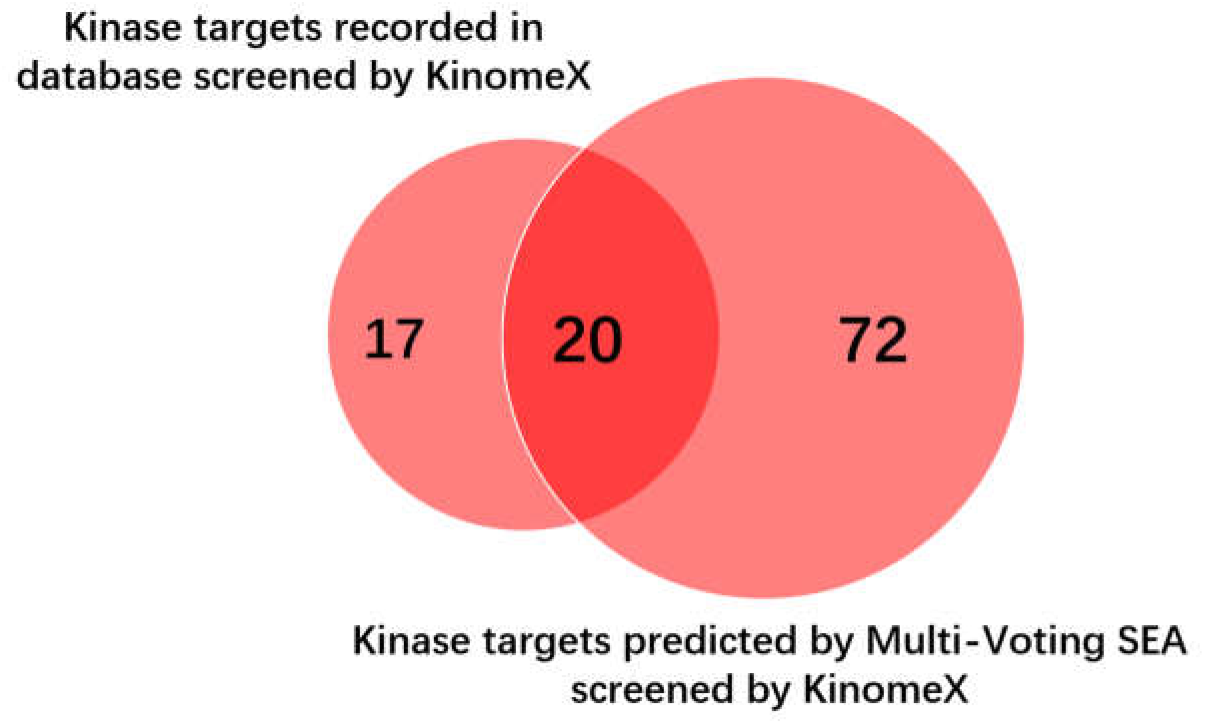
The number of potential direct kinase targets of CDDP to be verified

### 3 Active kinase targets of CDDP

Among the above 109 kinase targets, 106 kinase targets were contained in the commercial kinase spectrum developed by Eurofins Company. We test the activity of 106 kinase targets at 25 μg/ml concentration of CDDP. The active targets results were screened according to the threshold described in the method (Table 4).

**Table 4.** Statistical analysis on accuracy of activity test data for the potential direct kinase targets of CDDP

| Target sources | Targets screened by KinomeX | In Full KP | Active in Full KP | Accuracy (%) |
| --- | --- | --- | --- | --- |
| Recorded targets | 37 | 37 | 15 | 40.5 |
| Multi-Voting SEA predicted targets | 92 | 89 | 26 | 29.2 |
| Common targets | 20 | 20 | 11 | 55 |
| Total targets | 109 | 106 | 30 | 28.3 |

In total, 30 active targets were obtained, the accuracy was about 28.3% (Tables 4 and 5, Table S7). Among them, 15 out of 37 recorded kinase targets were verified, and the accuracy is about 40.5% (Table 4, Table S5). 26 out of 89 kinase targets predicted from Multi-voting SEA got active value, bringing the accuracy up to 29.2% (Table 4, Table S6). 11from common 20 kinase targets were verified, achieving 55% accuracy.

**Table 5.** Kinase target verified by activity test at 25 μg/ml concentration of CDDP

| # | Kinase | Protein | Gene | Gene ID | 25 µg/ml-Activity %* |
| --- | --- | --- | --- | --- | --- |
| 1 | Aurora-A(h) | Aurora-A | AURKA | 6790 | 76 |
| 2 | Aurora-B(h) | Aurora-B | AURKB | 9212 | 56 |
| 3 | Axl(h) | AXL | AXL | 558 | 73 |
| 4 | CaMKIIγ(h) | CaMK2gamma | CAMK2G | 818 | 64 |
| 5 | CLK1(h) | CLK1 | CLK1 | 1195 | 4 |
| 6 | Fms(h) | Fms | CSF1R | 1436 | 59 |
| 7 | CK1γ3(h) | CK1gamma3 | CSNK1G3 | 1456 | 58 |
| 8 | CK2α1(h) | CK2alpha1 | CSNK2A1 | 1457 | 65 |
| 9 | CK2α2(h) | CK2alpha2 | CSNK2A2 | 1459 | 64 |
| 10 | DYRK1A(h) | DYRK1A | DYRK1A | 1859 | 80 |
| 11 | FGFR1(h) | FGFR | FGFR1 | 2260 | 69 |
| 12 | Flt1(h) | Flt-1 | FLT1 | 2321 | 78 |
| 13 | Flt3(h) | FLT3 | FLT3 | 2322 | 74 |
| 14 | Fyn(h) | Fyn | FYN | 2534 | 63 |
| 15 | GSK3β(h) | GSK3beta | GSK3B | 2932 | 56 |
| 16 | HIPK4(h) | HIPK4 | HIPK4 | 147746 | 71 |
| 17 | IRAK4(h) | IRAK4 | IRAK4 | 51135 | 80 |
| 18 | Lck(h) | Lck | LCK | 3932 | 79 |
| 19 | MAPK1(h) | MAPK1(ERK2) | MAPK1 | 5594 | 78 |
| 20 | Met(h) | Met | MET | 4233 | 43 |
| 21 | TrkC(h) | TRKC | NTRK3 | 4916 | 72 |
| 22 | PDGFRα(h) | PDGFRalpha | PDGFRA | 5156 | 76 |
| 23 | Pim-1(h) | Pim1 | PIM1 | 5292 | 69 |
| 24 | Pim-2(h) | Pim2 | PIM2 | 11040 | 71 |
| 25 | Ret(h) | Ret | RET | 5979 | 61 |
| 26 | Rsk2(h) | RSK2 | RPS6KA3 | 6197 | 78 |
| 27 | SLK(h) | SLK | SLK | 9748 | 75 |
| 28 | SRPK1(h) | SRPK1 | SRPK1 | 6732 | 74 |
| 29 | DRAK1(h) | DRAK1 | STK17A | 9263 | 79 |
| 30 | Syk(h) | Syk | SYK | 6850 | 63 |

\* The lower the value, the stronger the binding activity.

Among them, 14 targets with active value lower than 70 were repeatedly verified at the concentration of 250 μg/ml, from which 9 targets with dose-dependent relationship were found, such as MET, PIM1 and SYK (Table 6).

**Table 6.** Comparison of the kinase targets verified by activity test at 25 μg/ml and 250 μg/ml of CDDP separately

| # | Kinase | Protein | Gene | Gene ID | 25 µg/ml-Activity %<sup>*</sup> | 250 µg/ml-Activity %<sup>*</sup> |
| --- | --- | --- | --- | --- | --- | --- |
| 1 | Aurora-B(h) | Aurora-B | AURKB | 9212 | 56 | 28 |
| 2 | CaMKIIγ(h) | CaMK2gamma | CAMK2G | 818 | 64 | 86 |
| 3 | CK1γ3(h) | CK1gamma3 | CSNK1G3 | 1456 | 58 | 23 |
| 4 | CK2α1(h) | CK2alpha1 | CSNK2A1 | 1457 | 65 | 47 |
| 5 | CK2α2(h) | CK2alpha2 | CSNK2A2 | 1459 | 64 | 43 |
| 6 | CLK1(h) | CLK1 | CLK1 | 1195 | 4 | 4 |
| 7 | FGFR1(h) | FGFR | FGFR1 | 2260 | 69 | 43 |
| 8 | Fms(h) | Fms | CSF1R | 1436 | 59 | 58 |
| 9 | Fyn(h) | Fyn | FYN | 2534 | 63 | 63 |
| 10 | GSK3β(h) | GSK3beta | GSK3B | 2932 | 56 | 39 |
| 11 | Met(h) | Met | MET | 4233 | 43 | 8 |
| 12 | Pim-1(h) | Pim1 | PIM1 | 5292 | 69 | 18 |
| 13 | Ret(h) | Ret | RET | 5979 | 61 | 70 |
| 14 | Syk(h) | Syk | SYK | 6850 | 63 | 2 |

\* The lower the value, the stronger the binding activity.

## Discussion

Aiming at the problems existing in the research of TCM prescriptions, this study considers that the key to solve the problem is to carry out the research on the direct targets of TCM prescriptions. In this study, we proposed a strategy based on recorded data with definite activity and algorithm prediction data to find the direct kinases target of TCM, which is independent of any specific disease model, and take CDDP as a case study. The results showed that we can quickly obtain the potential direct kinase targets at a lower cost and high reliability with high success rate of verification. The results can lay an important foundation for the study of the mechanism and material basis of TCM from the origin [10], expanding TCM potential new indications and promoting TCM modernization.

TCM prescriptions are the characteristics of TCM. These prescriptions embodies the dialectical thought of TCM and the medication holistic view. The ingredients contained in the prescriptions could be significantly different from those in the single herb. Therefore, in this study, the components contained in CDDP instead of those in single herb were taken as the research objects. As accumulating evidences have proved that the ingredients entering the blood, main metabolites, bioequivalence components compared to the prescription, and active components reported in literatures contribute more to the effects and mechanisms of TCM [35–37], we raised the hypothesis that the potential targets of all the important components mentioned above should be more likely to become the direct targets of the whole prescriptions. In addition, we included another component reported the most in single herbs but has not been confirmed in the whole prescription, quercetin as important component to finalize the list. This method undoubtedly improves the credibility of the data, which is different from most commonly used network pharmacology research flowchart [38–40]. In addition, in order to obtain the potential target data for the important components in the whole recipe, we integrated the recorded data and predicted data. On one hand, the existing research results have been fully utilized by comprehensively collecting the public activity data. On the other hand, in order to avoid missing some important targets, the algorithm model of target prediction based on structural similarity was used to predict the potential targets of components. Moreover, kinase targets predicted by KinomeX platform were used to filter the kinase targets obtained by the above two methods, which can further improve the success rate of further verification.

At present, there is no research on the direct target for any compound TCM as a whole. Most of the study focused on active components in TCM. In this study, potential targets of the important components in the complex system of TCM were taken as the start to carry out the direct targets of recipe in vitro. Among them, small molecule affinity chromatography and activity-based protein profiling (ABPP) are the most widely used target identification technologies for active ingredients extracted from Chinese herbal and other natural products. Target fishing technology is a widely used method, which is based on small molecule affinity chromatography [41] and the principle that drug molecules can covalently bind to proteins. The drug molecules are connected to biocompatible inert resin (magnetic beads or agarose gel) through chemical reaction. The target protein of the drug molecule was determined by electrophoresis, purification and high resolution mass spectrometry [42–46]. Using this strategy, a series of targets for active components of TCM have been successfully identified. For example, Tu research group constructed probe for sumitone [47] and chrysanthema lactone [48], successively identified the target of sumitone a as inosine monophosphate dehydrogenase 2 (IMPDH2), and the target of chrysanthema lactone with anti-inflammatory effect was heat shock protein 70 (HSP70). All of them revealed the mechanism of action for active components in TCM from the origin. The target fishing strategy is a highly maneuverable target identification technology, which effectively promotes the clarification for mechanism and speeds up the modernization process of TCM. In the process of using this strategy, if we need to introduce affinity tags with larger steric hindrance, the activity of the compounds may be reduced or even lost. Target fishing technology is suits more for further in-depth analysis to screen the specific ingredients binding individual targets.

14 targets with active value lower than 70 at the concentration of 25 μg/ml were verified again at the concentration of 250 μg/ml, from which 9 targets with dose-dependent relationship were found, such as MET, PIM1 and SYK. However, the activity of four kinases, CAMK2G, CSF1R, FYN and RET, increased. The possible reasons are as follows: firstly, the components with high molecular weight in TCM form great stereo-hindrance effect when the concentration increases, which hinders the combination between active molecules and targets. Secondly, due to the existence of positive effectors in CDDP, when the concentration increases, it produces a positive synergy through the allosteric effect, increasing the protein activity and relatively weakening the inhibition of the components on the activity of targets, such as the synergy weakens the affinity and internal effectiveness of the ligand on the receptors [49–53]. For example, the two components, CDDP 12 (rosmarinic acid) and CDDP 37 (catechol), contained in CDDP can act on the common target FYN. However, the binding site for the two components may be different, which may bring the allosteric effect, weakening the inhibition intensity under the condition of high concentration. These components are not directly binds to protein active sites, but the allosteric sites, outside the active sites of the protein, causing the conformational change of proteins and their activity. In the complex system of TCM, it may be due to the existence of allosteric effectors that the prescriptions can regulate the whole body in a systematic way.

In this study, 106 kinase targets were tested, and finally 30 active targets were obtained, with an accuracy of 28.3%. Among them, 15 out of 37 recorded kinase targets were verified, and the accuracy is about 40.5%. 26 out of 89 kinase targets predicted from Multi-voting SEA got active value, and the accuracy is about 29.2%. The 11 in the common 20 kinase targets were verified, the accuracy was 55%. As expected, the success rate of the known kinase targets is higher than that of the predicted targets. Hoverer, the candidate targets for verification is toolimited, and some key targets may be missed. By contrary, the targets obtained by predictive method can greatly expand the number of targets to be verified. The filter by the KinomeX predictive results enables a higher success rate at a lower cost. For any component of TCM prescription or western medicine, we can efficiently obtain the kinase targets to be verified through this system.

According to the current research results, the following related research can be carried out in the future: Firstly, we can predict and verify the specific components or component group in CDDP potentially regulating the active kinase targets obtained in this research. Thereby, the material basis of TCM can be elaborate. Secondly, we can use the active kinase targets to elucidate the mechanism of action for CDDP from a brand-new perspective. Thirdly, we can further compare and analyze the feature genes of some diseases and target genes modulated by CDDP, providing informative rationales for CDDP repositioning in the future.

The composition of CDDP is relatively simple, which only includs *Radix Salviae* (Danshen), *Panax Notoginseng (Burk.) F. H. Chen Ex C. Chow* (Sanqi), *Borneolum Syntheticum* (Bingpian). If we use the strategy mentioned above to screen direct targets for other complex prescriptions in the future, more factors will be taken into account Considering the complexity and diversity of ingredients in prescriptions, a variety of algorithms based on different principles to predict the relationship between components and targets can be used to obtain the potential target data of ingredients [54–58]. For example, the model predicting drug-target relationship based on network topology parameters [55], based on drug structure similarity and target structure similarity [56], based on clustering multi-dimensional drug target data [57], and based on deep learning and heterogeneous network [58] etc. By integrating the data predicted by various algorithms, the success rate may be improved. In addition, molecular docking technology can also be used to gain more reliable targets for further verification [59, 60].

## Conclusion

In this study, 30 direct targets of CDDP were obtained by the strategy mentioned above. This strategy is independent of any specific disease model, and can efficiently obtain the potential direct targets of TCM. Moreover, this method takes the TCM as a whole research object, which is in line with the holistic view and systematic theory of TCM, conforming to the guiding principles of pharmacology theory of TCM. The research results of the direct targets of TCM not only provide the theoretical basis for elucidating for the mechanism of action and the material basis, but also indicating rationales for the research of drug repositioning, which is of great significance for promoting TCM modernization.

## Supplemental information

Table S1 known targets of important ingredients obtained from public database.

Table S2 potential direct targets of important components predicted through Multi-voting SEA algorithm.

Table S3 potential direct targets of CDDP.

Table S4 protein kinase targets of important components predicted by KinomeX.

Table S5 37 kinase targets to be verified selected from known targets.

Table S6 92 kinase targets to be verified selected from targets predicted through Multi-voting SEA algorithm.

Table S7 potential direct kinase targets of CDDP to be verified.

Table S8 statistics of accuracy of activity test results for the target set to be verified without screening by targets obtained from KinomeX.

## Acknowledgments

The authors acknowledge the support provided by students in Professor Lin’s Lab.

## Declaration of interests

The authors declare no competing interests.

